# Frontotemporal dementia mutant tau (P301L) locks Fyn in an open, active conformation conducive to nanoclustering

**DOI:** 10.1101/2020.09.04.282855

**Authors:** Christopher Small, Ramón Martínez-Mármol, Tristan P. Wallis, Rachel S. Gormal, Jürgen Götz, Frédéric A. Meunier

## Abstract

Fyn is a Src kinase that controls critical signalling cascades and its postsynaptic enrichment underpins synaptotoxicity in Alzheimer’s disease (AD) and frontotemporal dementia (FTLD-tau). Previously, we found that pathogenic FTLD tau mutant (P301L) expression promotes aberrant trapping of Fyn in nanoclusters within hippocampal dendrites via an unknown mechanism (Padmanabhan et al., 2019). Here, we imaged Fyn-mEos2 using single particle tracking photoactivated localization microscopy (sptPALM) to demonstrate that nanoclustering of Fyn in hippocampal dendrites is promoted by Fyn’s open, primed conformation. Disrupting the auto-inhibitory, closed conformation of Fyn through phospho-inhibition, and perturbation of Fyn’s SH3 domain increases, Fyn’s nanoscale trapping. However, inhibition of Fyn’s catalytic domain has no impact on its mobility. Tau-P301L promotes Fyn lateral trapping via Fyn opening and ensuing increased catalytic activation. Pathogenic tau may therefore drive synaptotoxicity by locking Fyn in an open, catalytically active conformation, leading to postsynaptic entrapment and aberrant signalling cascades.

## Introduction

Fyn is a member of the Src family of kinases (SFKs), a group of enzymes that regulate signal transduction by catalysing the phosphorylation of tyrosine residues. Fyn is expressed in numerous cell-types, including lymphocytes, neurons and glia, and like other SFKs, is an intracellular, membrane-associated enzyme characterised by four conserved motifs known as Src homology domains (SH1, SH2, SH3 and SH4) (Figure 1A i). The SH1 domain is the catalytic domain responsible for the phosphorylation of tyrosine residues from target proteins and is connected via a polyproline type II (PPII) helix linker region to the SH2 domain (Figure 1A ii-iii). The SH2 and SH3 domains control the interaction of Fyn with its substrates, and the N-terminal SH4 domain is responsible for the association with the plasma membrane through myristylation and palmitoylation, thereby facilitating the interaction between Fyn and its membrane-associated substrates (Sato et al., 2009). The kinase activity of Fyn is also controlled by the transition between a closed (assembled) (Figure 1A ii) and an open (extended) conformation that can bind to its substrates and execute its catalytic activity (Figure 1Aiii). This transition is controlled by two conserved tyrosine phosphorylation sites with opposing roles. Dephosphorylation of Y531 in the C-terminal tail opens Fyn, resulting in its priming. This extended-primed form of Fyn is activated by trans-autophosphorylation of Y420, located in a central “activation loop” of the SH1 catalytic domain (Young et al., 2001). In the open conformation, the SH2 and SH3 domains are displaced and are free to interact with external ligands (Yadav & Miller, 2007). Conversely, two intramolecular interactions with the SH2 and SH3 domains are essential for downregulation of the kinase activity (Engen et al., 2008; Huculeci et al., 2016). Phosphorylation of Y531 allows this residue to interact with the SH2 domain and stabilises an assembled conformation whereby kinase activity is decreased by conformational changes at the active site of the catalytic domain. The second downregulatory interaction includes the SH3 domain and the PPII helix linker. Interestingly, this intramolecular interaction resembles the standard binding mode of SH3 domains to target sequences rich in proline and other hydrophobic amino acids (P-X-X-P motif, where P is proline and X is any amino acid). These sequences generally form a PPII helix which associates with the hydrophobic surface of the SH3 domain (Engen et al., 2008).

**Figure 1.**
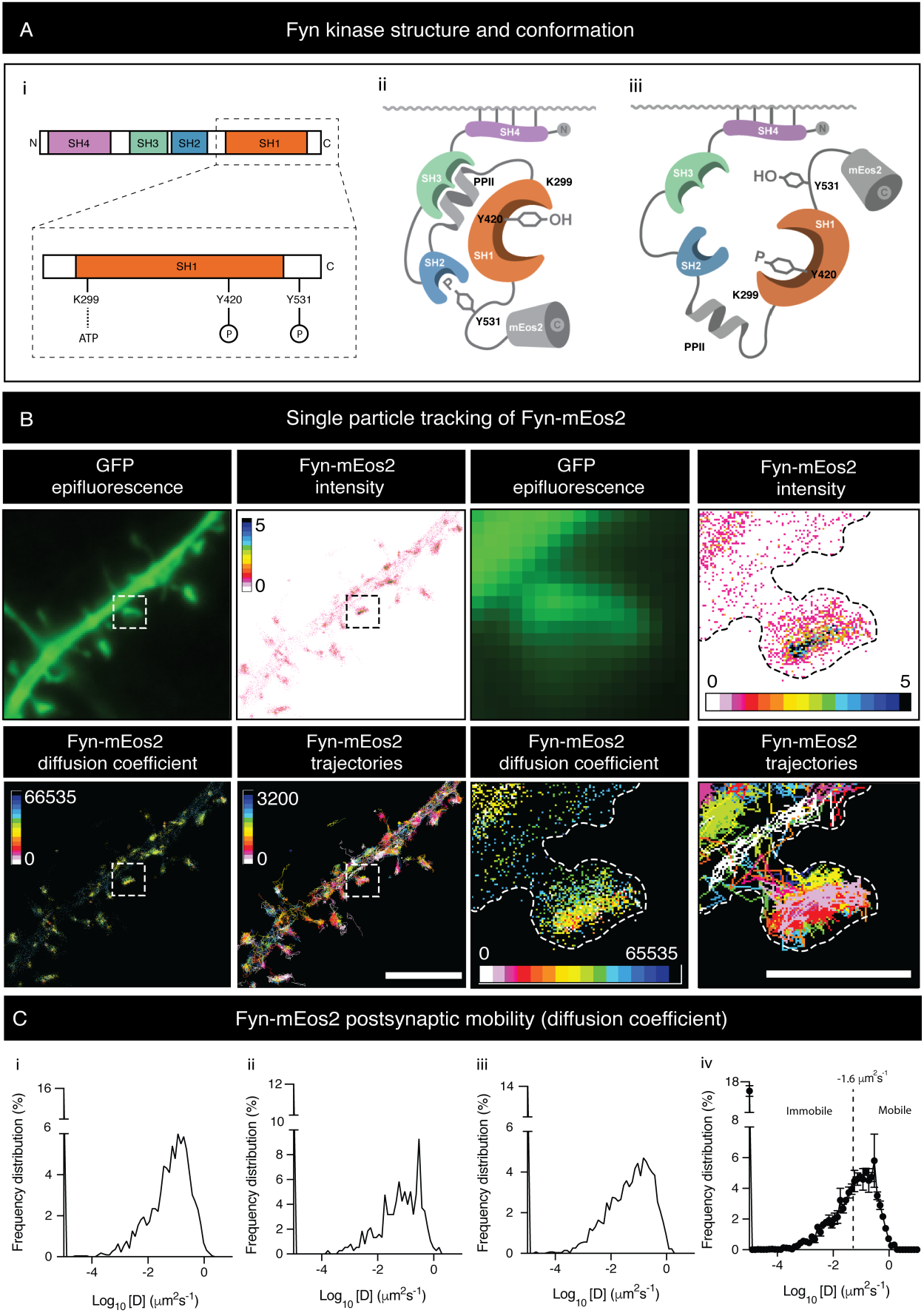
Single molecule imaging photoactivated localization microscopy (sptPALM) of Fyn-mEos2 co-transfected with GFP in secondary dendritic branches and spines of hippocampal neurons (DIV19-22). **A**. Illustration showing the structure and conformation of Fyn-mEos2. Key epitopes found in the SH1 domain and C terminus of Fyn are highlighted. **(i)** Secondary structure and domain alignment of Fyn. **(iii)** Tertiary structure of Fyn, closed conformation **(iii)** Tertiary structure of Fyn, open conformation. mEos2 is conjugated to the C-terminus of Fyn. **B**. Panels depict GFP epifluorescence and intensity, diffusion coefficient and tracking maps of Fyn-mEos2 (note: hotter colours within diffusion coefficient maps designate regions of low mobility). Scale bar = 4 µm (dendrites) and 1 µm (spines). **C**. Frequency distribution (%) of postsynaptic Fyn-mEos2 mobility, Log_10_[D](µm^2^s^-1^), where [D] is the diffusion coefficient. **(i-iii)** Examples of frequency of diffusion of Fyn-mEos2 from individual neurons. **(iv)** Averaged frequency distribution. Mobile:immobile fraction threshold is set at -1.6 µm^2^ s^-1^.

Fyn is responsible for integrating multiple signalling cascades. In neurons, Fyn facilitates cell-to-cell communication by promoting the scaffolding of the N-methyl-D-aspartate (NMDA) receptor through phosphorylation of the NR2B subunit of the receptor at the Y1472 epitope. This increases the affinity of NR2B with postsynaptic density protein 95 (PSD95) and, as a consequence, facilitates the stabilisation of NMDA receptor clusters at the membrane, thereby maintaining postsynaptic excitatory currents (Tezuka et al., 1999). Fyn is also implicated in mediating neurodegeneration. Overactive Fyn is believed to exacerbate cell death by promoting excess activation of NMDA receptors, leading to aberrant calcium entry into the postsynapse and neuronal excitotoxicity (Xia & Gotz, 2014). Fyn has also been shown to play a critical role in the neurotoxicity of Alzheimer’s disease (AD) and frontal temporal lobar degeneration with tau (FTLD-tau) (Briner et al., 2020; Polanco et al., 2018). Fyn promotes neurotoxicity downstream of the amyloid-beta (Aβ) peptide, which forms extracellular amyloid plaques, and the microtubule-associated protein tau (MAPT). Hyperphosphorylated tau accumulates into neurofibrillary tangles (NFTs), a key hallmark of the toxicity associated with AD and FTLD-tau (Gotz et al., 2001). In transgenic mouse lines that replicate AD pathology, knockout of Fyn has been shown to protect against the loss of presynaptic terminals and to delay premature mortality (Chin et al., 2004). Hippocampal slices from Fyn knockout mice have been shown to resist the toxic effects of treatment with toxic Aβ oligomers (Um et al., 2012). Similarly, pharmacological inhibition of Fyn rescues the behavioural deficits observed in AD mice (Kaufman et al., 2015). Conversely, overexpression of Fyn exacerbated the neuronal deficits present in AD mice (Chin et al., 2005; Kaufman et al., 2015).

Increasing evidence suggests that Fyn works as a key partner in tau-mediated pathology. Fyn interacts with tau through its SH2 and SH3 domains and P-X-X-P motifs located in the proline-rich region of tau (Bhaskar et al., 2005; Lau et al., 2016; Lee et al., 1998). This interaction mediates phosphorylation of tau at residue Y18, one of the epitopes associated with the formation of NFTs in AD patients (Bhaskar et al., 2010; Lee et al., 2004; Miyamoto et al., 2017; Neddens et al., 2018), and potentiates its association with Fyn (Usardi et al., 2011). Subcellular compartmentalisation of Fyn and tau also appears to be crucial for their toxicity. Mis-localisation of tau into dendritic spines mediates the synaptic dysfunction associated with AD and FTLD-tau (Frandemiche et al., 2014; Hoover et al., 2010; Miller et al., 2014; Xia et al., 2015). Fyn regulates the phosphorylation and dendritic distribution of tau through its activation by Aβ during AD (Larson et al., 2012). Alternatively, a previous study found that tau itself controls the localisation of Fyn to dendrites, and that disruption of postsynaptic targeting of Fyn mitigates Aβ toxicity (Ittner et al., 2010).

Altered organisation of receptors and signalling molecules in nanodomains is emerging as a key regulator of neuronal toxicity (Shrivastava et al., 2017). Several proteins involved in AD show aberrant clustering associated with the pathology. The metabotropic glutamate receptor mGluR5, for example, shows increased clustering when binding to Aβ (Renner et al., 2010); and the amyloid precursor protein (APP) organises into regulatory nanodomains that control the availability of APP molecules for proteolytic processing (Kedia et al., 2020). In this respect, tau has been shown to control the lateral trapping of Fyn into nanodomains within the dendrites and spines of hippocampal neurons, with single molecule imaging of Fyn-mEos2 showing enhanced mobility and decreased clustering in tau knockout neurons. Importantly, expression of an FTLD tau mutant (P301L) aberrantly increases the number of Fyn nanoclusters in spines, an effect that is likely to contribute to NMDA-mediated excitotoxicity (Padmanabhan et al., 2019). However, whether the nanoscale spatiotemporal organisation of Fyn is involved in its efficient transactivation remained to be established, and the mechanisms underlying the Fyn-tau toxic partnership have not yet been fully elucidated.

In this study, we used single-particle tracking photoactivated localization microscopy (sptPALM) to determine how the nanoscale organisation of Fyn is affected by its activity and conformation in the dendrites of live hippocampal neurons. Through pharmacological inhibition, blocking phosphorylation of the Y420 residue, preventing binding of ATP to the K299 epitope, or introducing a phosphorylation-inhibitory mutant to render Fyn constitutively open, we evaluated whether preventing the catalytic activity or altering the overall conformation can modulate Fyn mobility dynamics in neurons. Our results demonstrate that the SH3 domain is essential for maintaining a closed conformation and interacting with Fyn-binding proteins such as the FTLD-associated P301L mutant tau. Fyn entry into an opened, primed conformation is associated with its nanoscale entrapment. This extended and confined molecular configuration is compatible with the binding of pathological mutant tau to Fyn. Taken together, our findings suggest a molecular mechanism in which binding of P301L tau to Fyn in the dendritic compartment exacerbates Fyn’s toxic effects in FTLD-tau.

## Results

### The catalytic activity of Fyn does not alter the nanoscale organisation of Fyn in hippocampal dendrites

We recently discovered that Fyn displays a nanocluster organisation in the dendritic spines of hippocampal neurons that is dynamically regulated by neuronal maturation (Padmanabhan et al., 2019). In this study, we investigate whether changes in Fyn activity affect the nanoscale organisation of Fyn in live neurons. Fyn-mEos2 was expressed in hippocampal neurons, together with mCardinal or GFP as a cytoplasmic marker, and we performed sptPALM in an oblique illumination configuration. We imaged mature neurons at DIV19-22 and selected secondary dendrites with mature spines for analysis (Figure 1B). Fyn-mEos2 molecules were randomly photoconverted from a green-to a red-emitting state in response to constant weak illumination at 405 nm at a low spatial density. The photoconverted molecules were tracked in the red-emitting channel at 561 nm excitation at 50 Hz for a duration of 320 s (16,000 frames), in order to resolve the nanoscale distribution and dynamics of Fyn at a high spatiotemporal resolution in live neurons (Figure 1C).

Given that Fyn has a critical role in integrating a multitude of signalling pathways in neurons (Li & Gotz, 2017), we first sought to determine whether its catalytic activity was involved in promoting its nanoclustering at the post-synapse. Based on experiments performed with other SFKs, it is established that mutations of specific residues in the SH1 domain have profound effects on the activity of these enzymes, without affecting their conformation (Engen et al., 2008; Nika et al., 2010; Sicheri & Kuriyan, 1997). Trans-autophosphorylation of Y420 located at the central “activation loop” of the catalytic domain is required for the transition from an open, primed to an opened, active form able to phosphorylate its substrates (Nika et al., 2010; Young et al., 2001). To determine if the trans-autophosphorylation of Y420 in the catalytic SH1 domain controls the nanoclustering of Fyn, we introduced a phospho-inhibitory mutation (Y420F) to generate a ‘kinase-inactivated’ Fyn enzyme (Figure 2A-B). Our results showed that this mutation did not affect the mobility of Fyn in dendritic branches of hippocampal neurons (Figure 2C i-ii). Similarly, no changes in the mobility of Fyn-mEos2 were observed in dendritic spines (Figure 2D i-ii). To further evaluate the importance of Fyn activity, we generated a ‘kinase-dead’ Fyn by introducing the mutation K299M, designed to block interaction with ATP, thereby creating an inactive enzyme unable to phosphorylate other substrates (Figure 2A) (Jin et al., 2017; Twamley et al., 1992; Twamley-Stein et al., 1993). Similarly, the Fyn-K299M-mEos2 mutation had no effect on the mobility compared to wild-type Fyn-mEos2 in dendrites (Figure 2C i-ii) and spines (Figure 2D i-ii). Blockade of Fyn activity was also achieved by incubating neurons with 3-(4-chlorophenyl)-1-(1,1-dimethylethyl)-1H-pyrazolo[3,4-d]pyrimidin-4-amine (PP2) (10 μM, 30 minutes), a potent and specific pharmacological inhibitor of SFK’s kinase catalytic activity (Hanke et al., 1995; Jin et al., 2017; Tomatis et al., 2013). We used the structurally related inactive analogue 1-phenyl-1H-pyrazolo[3,4-d]pyrimidin-4-amine (PP3) (10 μM) as a control that does not inhibit Src family members (Figure 2 – Figure 1 supplement) (Bain et al., 2003). PP2 is an ATP-competitive inhibitor that interacts with the hydrophobic pocket near the ATP-binding cleft of the SH1 domain, thereby preventing the binding of the substrate ATP to the enzyme (Zhu et al., 1999). No alterations in Fyn-mEos2 mobility were observed in the dendrites of hippocampal neurons in response to PP2 or PP3 treatment (Figure 2-Figure 1 supplement) suggesting that the catalytic activity had no influence on the nanoscale organisation of Fyn at the synapse. Overall, these results further support the notion that the catalytic activity of Fyn does not influence its nanoscale organisation at the post-synapse.

**Figure 2.**
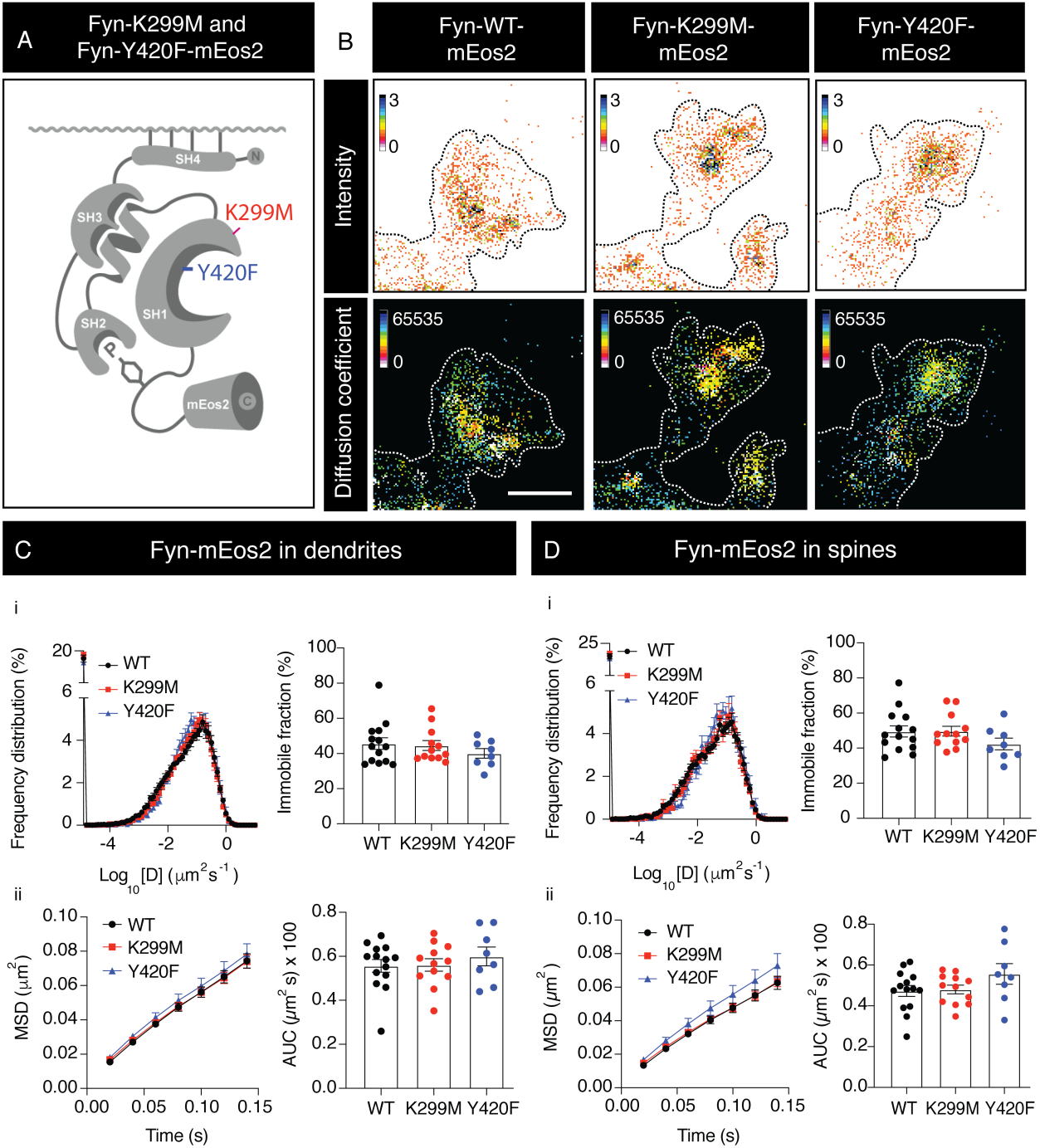
Inhibition of the catalytic (SH1) domain does not impact the mobility of Fyn-mEos2 **A**. Illustration depicting K299M and Y420F mutants in closed conformation of Fyn. **B**. Intensity and diffusion coefficient maps for Fyn-mEos2 (wild-type, K299M, Y420F). **C**. Mobility of Fyn-mEos2 in dendrites. **D**. Mobility of Fyn-mEos2 in spines. **(i)** Frequency distribution of Log_10_ [D](µm^2^s^-1^) where [D] is the diffusion coefficient, together with the immobile fraction (%). **(ii)** Mean square displacement (MSD) of Fyn-mEos2 over time (0.14 sec) with corresponding area under curve (AUC) [(µm^2^ s) × 100]. Error bars are standard errors of the mean (SEM). Mean ± SEM values were obtained for neurons transfected with Fyn-WT-mEos2 (n = 14), Fyn-K299M-mEos2 (n=12) and Fyn-Y420F-mEos2 (n=8). Statistical comparisons were performed using a Student’s t test.

### Induction to an open, primed conformation promotes the nanoscale entrapment of Fyn in hippocampal dendrites and spines

The closed conformation of Fyn is primarily maintained through phosphorylation of the Y531 epitope in the C-terminal tail, which stabilises the interaction of the C-terminal domain of Fyn with its SH2 region. To constitutively force an open conformation, we introduced a phospho-inhibitory mutation (Y531F) into the C-terminal domain of Fyn (Figure 3A). The opened-primed conformation rapidly facilitates the trans-autophosphorylation on Y420 that results in a fully active enzyme. In the presence of the Y531F mutation, Fyn is unable to return to its inactive-closed conformation, resulting in a ‘constitutively-active’ form of Fyn (Nakazawa et al., 2001; Xia & Gotz, 2014). To examine the effect of Y531F-induced constitutively open conformation on the mobility of Fyn in dendrites, we performed sptPALM of Fyn-Y531F-mEos2. We observed a significant decrease in Fyn mobility, indicating that the open conformation of Fyn is essential for mediating the nanoscale lateral trapping of Fyn within dendrites (Figure 3C i-ii) and spines (Figure 3D i-ii). To evaluate whether the kinase activity of Fyn played a role in its immobilization, or whether this effect was solely associated with a change in the conformation towards an extended one, we created the mutant Fyn-Y531F-K299M, that remains constitutively opened but with an inactive catalytic domain (Figure 3A). This double mutant also showed significantly lower mobility than the wild-type Fyn in dendrites (Figure 3C i-ii) and spines (Figure 3D i-ii). Interestingly, our results revealed an attenuation of the opened Fyn immobilization in the presence of the K299M mutation (Figure 3C-D). This observation suggests that modifying the catalytic domain through the K299M mutation somehow interferes with Fyn’s open conformation. In summary, these results demonstrate that the nanoscale organisation of Fyn is principally controlled by its entry into an open, primed conformation through dephosphorylation of the Y531 epitope.

**Figure 3.**
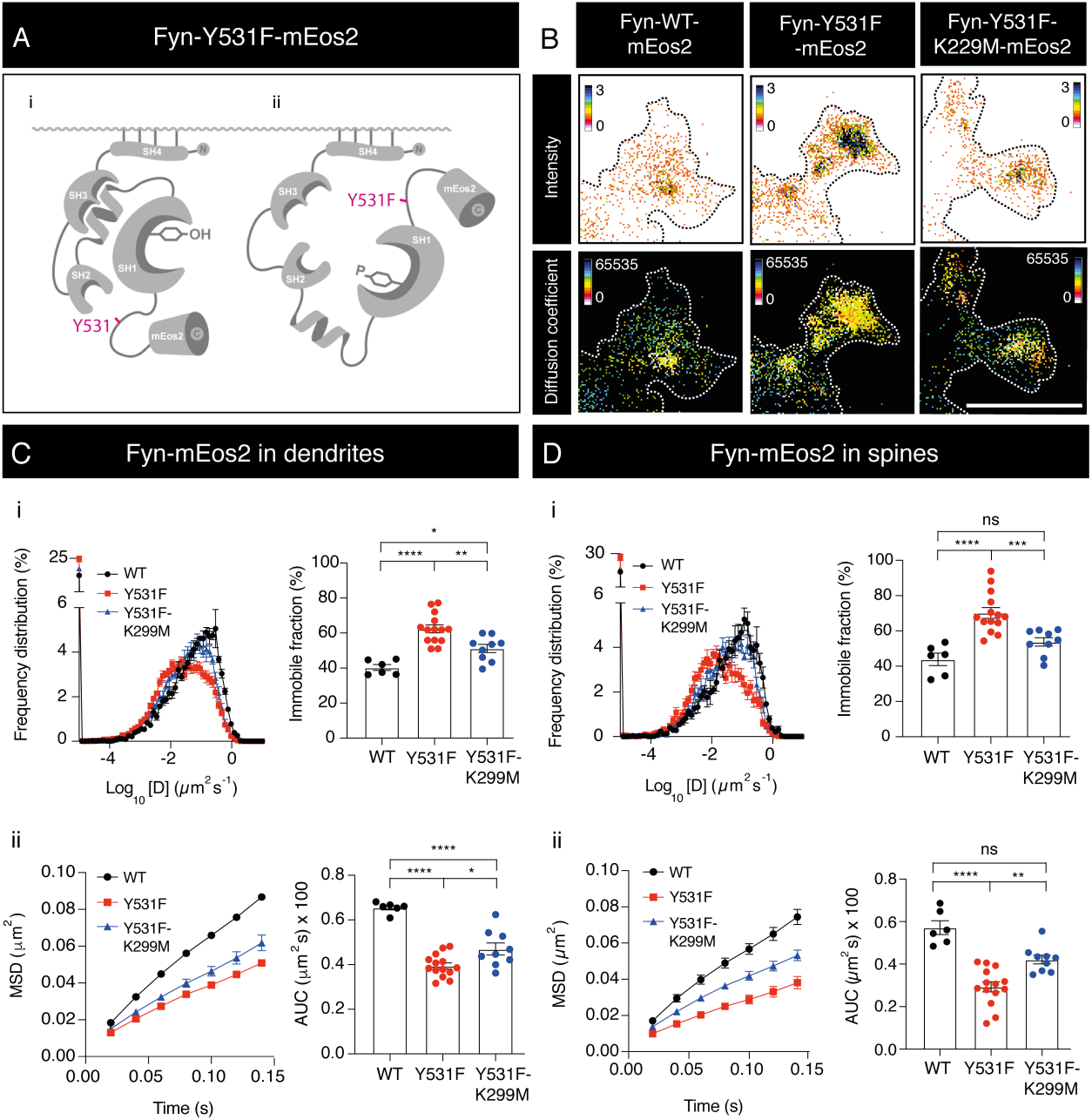
Phosphorylation of the Y531 epitope controls the lateral entrapment of Fyn-mEos2 in the dendrites and spines of hippocampal neurons. **A**. Illustration depicting **(i)** the Y531 epitope in the closed conformation of Fyn and **(ii)** the Y531F mutation triggering an open conformation of Fyn. **B**. Intensity and diffusion coefficient maps of Fyn-mEos2 localisations within spines of hippocampal dendrites. Scale bar = 1 µm. **C**. Mobility of Fyn-mEos2 in hippocampal dendrites. **(i)** Frequency distribution of diffusion coefficient values together with immobile fraction (%). **(ii)** Mean square displacement (MSD) of Fyn-mEos2 over time (0.14 s) with the corresponding area under curve (AUC) [(µm^2^ s) × 100]. **D**. Mobility of Fyn-mEos2 in hippocampal spines. **(i)** Frequency distribution of diffusion coefficient values together with immobile fraction (%). **(ii)** Mean square displacement (MSD) of Fyn-mEos2 over time (0.14 s) with the corresponding area under curve (AUC) [(µm^2^ s) × 100]. Error bars are standard errors of the mean (SEM). Mean ± SEM values were obtained for neurons transfected with Fyn-WT-mEos2 (n = 8), Fyn-Y531F-mEos2 (n=14) and Y531F-K299M-mEos2 (n=9). Statistical comparisons were performed using a one-way ANOVA and Tukey’s test for comparisons between groups.

To determine if the changes in mobility observed with the Y531F mutant were a product of an alteration in the lateral trapping of Fyn-mEos2 molecules, we compared the frequency distribution (%) of inter-frame step lengths taken by Fyn-WT-mEos2 and Fyn-Y531F-mEos2 throughout our acquisitions. Interestingly, the average inter-frame step size of Fyn-WT-mEos2 throughout its lifespan was not significantly different from that of the Y531F mutant (Figure 4A), suggesting that the changes in mobility of the Y531F mutant may be due a perturbation in the nanoclustering of Fyn-mEos2. To investigate this, we used Density-Based Spatial Clustering of Applications with Noise (DBSCAN) to quantify the size and density of Fyn-mEos2 nanoclusters (Figure 4B-D). This approach involved evaluating the spatial distribution of Fyn-mEos2 trajectory centroids, which allowed us to determine which of those trajectories were confined within spatial clusters, and to derive metrics on cluster number, area, density and trajectory mobility. DBSCAN analysis confirmed that wild-type Fyn had decreased mobility when organised in small nanoclusters at the synapse (Figure 4B) that occupy an average area of 0.271 ±0.016 μm^2^. Within these nanoclusters, the mobility and displacement of Fyn were restricted (Figure 4C i). The Y531F mutation, which induces an active, open conformation, caused Fyn-mEos2 to have a significantly lower mobility (MSD) within nanoclusters (Figure 4C ii). Furthermore, the Y531F mutation caused Fyn-mEos2 to be packaged into smaller nanoclusters (0.124 ± 0.006 μm^2^) with increased trajectory membership (Figure 4D i) and of significantly higher density (Figure 4D ii-iii). These results led us to conclude that the decreased mobility of Fyn-mEos2 observed in its open conformation was due to increased nanoclustering propensity, rather than a decrease in the size of the inter-frame steps of Fyn-mEos2 molecules.

**Figure 4.**
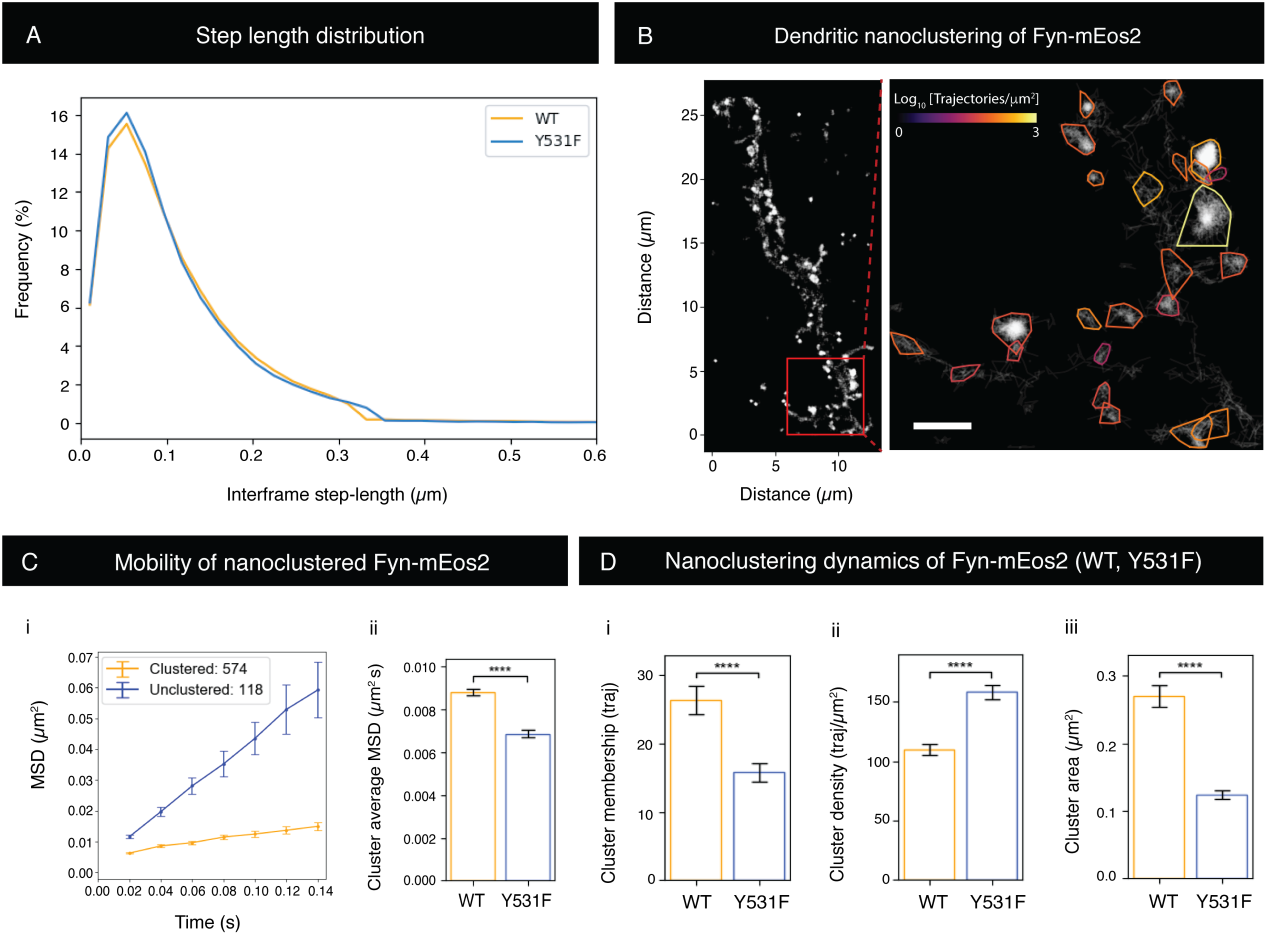
Nanoclustering and step length distribution of Fyn-mEos2 in the dendrites of hippocampal neurons revealed by density-based clustering analysis (DBSCAN). The Y531F mutation, which induces entry of Fyn into an open, active conformation, increases the density and decreases the size of Fyn-mEos2 nanoclusters. **A**. Frequency distribution of step lengths (µm) of Fyn-mEos2. No change in the frequency distribution of steps taken was observed between Fyn-WT-mEos2 and Fyn-Y531F-mEos2. **B**. Spatial distribution of Fyn-mEos2 nanoclusters in the dendrites and spines. Scale bar = 1 µm. **C. (i)** Mean square displacement (MSD) (µm^2^) over time (200 ms) plotted for clustered and unclustered populations of Fyn-mEos2. **(ii)** Average cluster MSD AUC (µm^2^s) for Fyn-WT-mEos2 and Fyn-Y531F-mEos2. **D**. Nanocluster dynamics of Fyn-WT-mEos2 and Fyn-Y531F-mEos2. (**i)** Cluster membership (trajectory number). (**ii)** Cluster density (trajectories/µm^2^). (**iii)** Cluster area (µm^2^). Nanoclusters were significantly denser for Fyn-Y531F-mEos2 compared to Fyn-WT-mEos2. Error bars are standard errors of the mean (SEM). Mean ± SEM values were obtained from n = 823 nanoclusters (Fyn-WT-mEos2) and n=698 (Fyn-Y531F-mEos2) from 5 neurons. Statistical comparisons were performed using a Student’s t test.

### Alteration of the SH3 domain renders Fyn more immobile in dendrites

The intramolecular interaction between the SH3 domain and the PPII helix linker is also essential to stabilise the inactive, closed conformation of Fyn and other SFKs (Moroco et al., 2014; Young et al., 2001). As a consequence, displacement of the SH3 domain from the helix linker has been reported to facilitate the activation of these kinases (Briggs & Smithgall, 1999; Moroco et al., 2014). The SH3 domain is also involved in the interaction of Fyn with its substrates such as the heterogeneous nuclear ribonucleoprotein A2 (hnRNPA2) (Amaya et al., 2018), p85*α* (Morton et al., 1996), the palmitoyl-acyl transferase DHHC5 (Brigidi et al., 2015), tau (Lee et al., 1998; Reynolds et al., 2008) and P301L mutant tau (Bhaskar et al., 2005). As displacement of the SH3 domain has been used to activate Fyn molecules and other SFKs (Moroco et al., 2014), we wanted to explore whether direct alterations of this domain affects the nanoscale organisation of Fyn. To investigate this, we performed sptPALM on a Fyn protein lacking the SH3 domain (Fyn-ΔSH3-mEos2) (Figure 5A, B). Deletion of the SH3 domain resulted in a decreased mobility of Fyn (Figure 5C, D). In accordance to what has been observed through alteration of the SH3 domain in other SFKs (Moroco et al., 2014), our results may suggest that the absence of the SH3 domain destabilises the closed conformation of Fyn. Similar to the Y531F mutation, this could facilitate the transition of Fyn to the open, primed conformation, leading to Fyn’s immobilisation.

**Figure 5.**
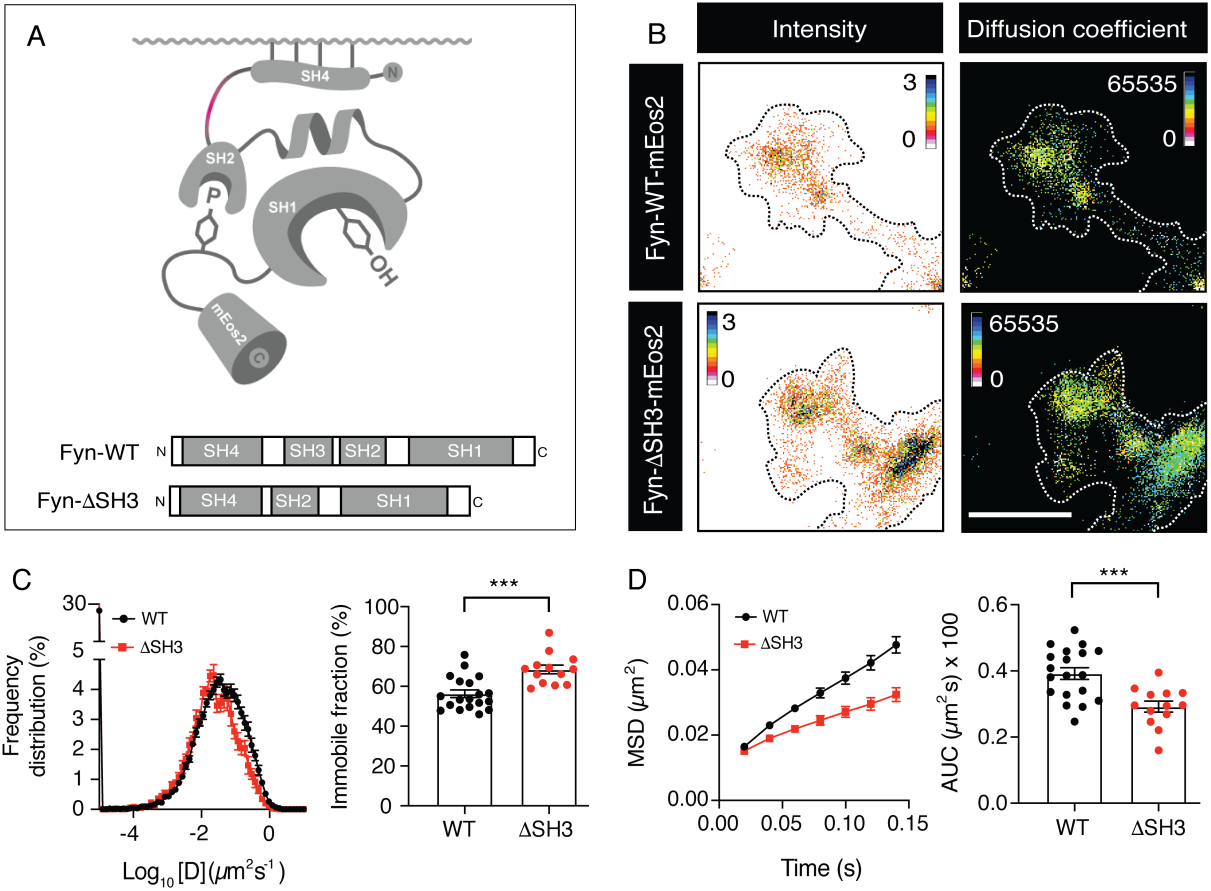
Deletion of the SH3 domain promotes the lateral trapping of Fyn-mEos2. **A**. Schematic showing deletion of the SH3 domain (ΔSH3). **B**. Intensity and diffusion coefficient maps of Fyn-mEos2 (WT, ΔSH3) in hippocampal neurons. Scale bar = 1 µm. **C**. Frequency distribution of diffusion coefficient values, where [D]=diffusion coefficient, together with the immobile fraction (%). **D**. Mean square displacement (MSD) over time (0.14 s) with corresponding area under curve (AUC) [(µm^2^ s) × 100] of Fyn-mEos2 (WT, ΔSH3) in hippocampal dendrites (DIV18-22). Error bars are standard errors of the mean (SEM). Mean ± SEM values were obtained from neurons transfected with Fyn-WT-mEos2 (n=19) and Fyn-ΔSH3-mEos2 (n = 13). Statistical comparisons were performed using a one-way ANOVA and Tukey’s test for comparisons between groups.

Due to the importance of the SH3:helix linker interaction in stabilising the closed conformation of SFKs, several strategies have been developed to manipulate the activity of these enzymes based on the creation of small molecules that selectively bind the SH3 domain (Huang et al., 2016; Kukenshoner et al., 2017; Moroco et al., 2014; Yadav & Miller, 2007). The G9 monobody was specifically generated to interact with the SH3 domain, displaying high specificity for Fyn among other kinases (Huang et al., 2012). To further evaluate how alterations of the SH3-PPII helix linker interaction affects the nanoscale organisation of Fyn, we created a mEos2 tagged version of the G9 monobody (Figure 6A). As a control, we used a mEos2-tagged anti-GFP nanobody co-transfected with Fyn-GFP in HEK293 cells (Figure 6A, B). Our results demonstrate that the anti-GFP-mEos2 nanobody, co-expressed with Fyn-GFP, displays similar mobility to Fyn-mEos2 alone. However, co-expression of Fyn-GFP together with the mEos2 G9 anti-Fyn-mEos2 monobodies targeting the SH3 domain, resulted in a significant decrease in the MSD of Fyn-GFP in HEK293T cells (Figure 6C). This result suggests that binding of monobodies to the SH3 domain of Fyn causes an increase in the immobilisation of Fyn molecules. Interfering with the SH3-PPII helix linker region may therefore facilitate the acquisition of a conformation conducive to lateral trapping of extended-primed Fyn molecules.

**Figure 6.**
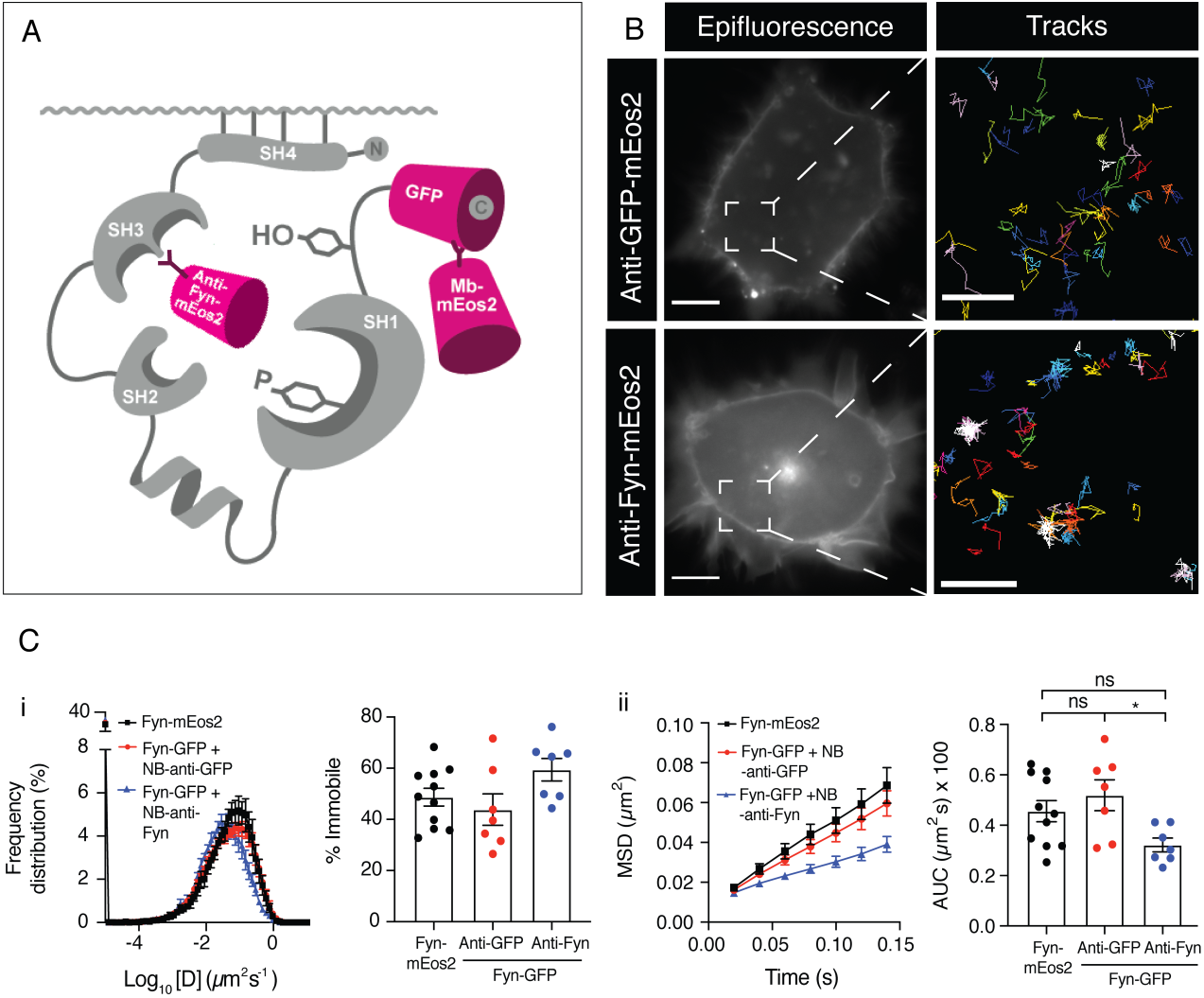
Binding of anti-Fyn intrabodies to the SH3 domain decreases the mobility of Fyn-GFP in HEK293T cells. **A**. Schematic depicting binding of anti-Fyn-mEos2 intrabodies to the SH3 domain and anti-GFP-mEos2 intrabodies to Fyn-GFP. **B**. Anti-GFP-mEos2 and anti-Fyn (SH3)-mEos2 tracking of Fyn-GFP in HEK293T cells. Scale bar = 4 µm for epifluorescence and 1 µm for track panels. **C. (i)** Frequency distribution of diffusion coefficient values together with immobile fraction (%) and **(ii)** MSD over time (0.14 sec) with AUC [(µm^2^ s) × 100] of Fyn-mEos2, anti-GFP-mEos2/Fyn-GFP or anti-SH3-mEos2/Fyn-GFP in HEK293T cells. Error bars are standard errors of the mean (SEM). Mean ± SEM values were obtained from HEK293T cells transfected with Fyn-mEos2 (n=11), Fyn-GFP treated with anti-GFP-mEos2 (n=7) or anti-Fyn-mEos2 (n=7). Statistical comparisons were performed using a one-way ANOVA and Tukey’s test.

### Overexpression of wild-type and FTLD-tau mutant (P301L) tau promotes an open conformation and immobilisation of Fyn in hippocampal dendrites

Previous findings suggest that aberrant nanoclustering is enhanced by the presence of a FTLD mutant form of tau (P301L) (Padmanabhan et al., 2019). This mutant tau binds to the SH3 domain of Fyn with higher affinity than wild-type tau (Bhaskar et al., 2005). To further evaluate the effect of wild-type and P301L mutant tau on the mobility of Fyn-mEos2, we co-expressed wild-type tau and P301L mutant tau with Fyn-mEos2 in hippocampal neurons and performed sptPALM imaging (Figure 7A). Wild-type tau caused a significant decrease in the mobility of Fyn-mEos2 (Figure 7B, C). In accordance with our previous findings, this decrease was accentuated in the presence of P301L tau (Figure 7B, C), which has an increased binding capacity to Fyn (Bhaskar et al., 2005). This is interesting as tau binds Fyn through its SH3 domain, which causes a decrease in Fyn mobility when disrupted. These results suggest that the intramolecular interaction between the SH3 and PPII helix linkers regions plays a key role in controlling the nanoscale organisation of Fyn, and that wild-type and P301L mutant tau drives the nanoclustering of Fyn by displacing this interaction and thus stabilising an open and highly immobile Fyn conformation. We co-expressed Fyn and tau or tau-P301L in HEK293T cells and observed that the total expression of Fyn increased in the presence of P301L mutant tau, and that the proportion of phosphorylated Fyn (Y420) was also higher (Figure 7D). Taken together, these findings suggest that the interaction of Fyn with P301L tau exposes and primes the Y420 epitope for phosphorylation, a process which is associated with increased nanoclustering in hippocampal neurons.

**Figure 7.**
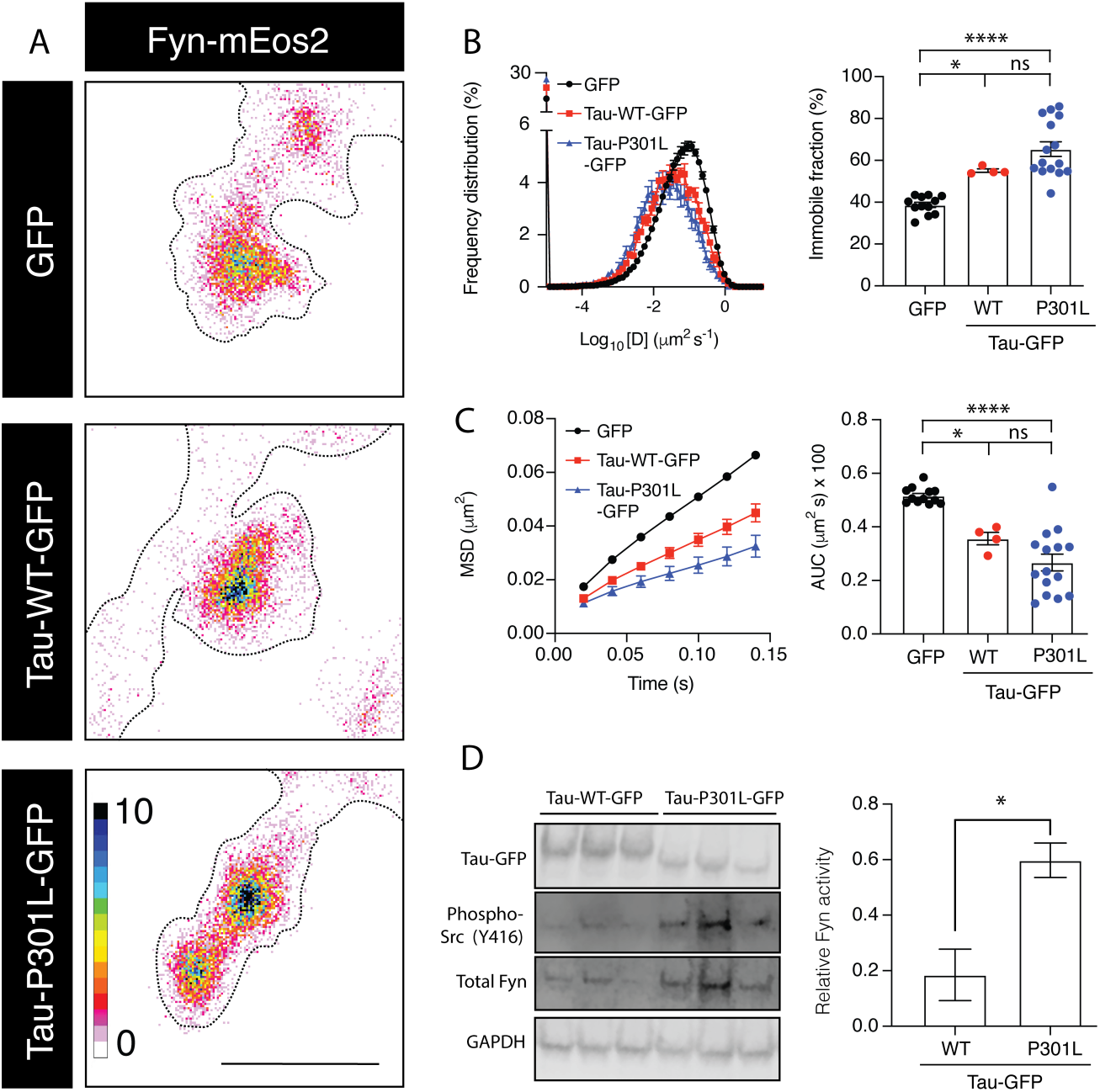
Overexpression of tau (WT, P301L) decreases the mobility and increases activity of Fyn. **A**. Intensity map of Fyn-mEos2 co-expressed with GFP, tau-WT-GFP and tau-P301L-GFP in the spines of mouse hippocampal neurons (DIV19-22). Scale bar = 1 µm. **B**. Frequency distribution (%) of Log_10_ [D] of Fyn-mEos2 with immobile fraction (%). **C**. Mean square displacement (MSD) of Fyn-mEos2 over time (0.14 sec) and area under curve (AUC) [(µm^2^ s) × 100]. **D**. Increased phosphorylation at the Y16 epitope in the presence of tau P301L. Western blot of Fyn in protein extracts derived from HEK293 cells co-transfected with Fyn-WT-mEos2 and GFP, tau-WT-GFP or tau-P301L-GFP. Fyn-Y420F-mEos2 was also co-transfected with GFP as a control. Activity of Fyn determined by quantifying proportion of active Fyn (anti-Y416) relative to total Fyn, normalised to GAPDH. Co-expression of tau-P301L increased the proportion of active Fyn. Error bars are standard errors of the mean (SEM). Mean ± SEM values were obtained from neurons transfected with Fyn-mEos2+GFP (n = 12), Fyn-mEos2+tau-WT-GFP (n=4) or Fyn-mEos2+tau-P301L-GFP (n=15), and HEK293 protein lysates (n=3 for each group). Statistical comparisons between tau-transfections (A-C) were performed using a one-way ANOVA and Tukey’s test for multiple comparisons. A Student’s t test was performed for statistical comparisons of Y16 phosphorylation (D).

## Discussion

Fyn is an SFK whose activity integrates multiple signalling cascades. However, when dysregulated, Fyn has been associated with the development of neurological disorders such as AD and tau pathologies (Haass & Mandelkow, 2010; Ittner & Gotz, 2011). We recently provided the first conceptual framework for how the nanoscale organisation of Fyn is altered during tau-mediated neurodegeneration (Padmanabhan et al., 2019). However, it is not known whether the nanoscale spatiotemporal organization of Fyn is required for its efficient signal transduction, and the mechanisms driving the toxic partnership of Fyn and tau remains unsolved. Our new results now suggest that the transition of Fyn into an open conformation facilitates its organization into nanoclusters, and that the binding of toxic P301L mutant tau associated with FTLD-tau locks Fyn into an open, immobile and catalytically active conformation, thereby providing a rational explanation for the resulting initiation of aberrant toxic signalling cascades.

The kinase activity of Fyn and other SFKs is controlled by the transition between two opposing configurations (from closed/inactive to open/primed). This transition allows SFKs to bind their substrates and execute their catalytic activity (Sicheri & Kuriyan, 1997). Closed Fyn is locked through two intramolecular interactions between the SH2 domain bound to phosphorylated Y531 at the C-terminal tail, and the interaction between the hydrophobic residues from the SH3 domain with the proline-rich PPII helix linker (Engen et al., 2008). Our results showed that altering any of the locking interactions reduces the mobility of Fyn, suggesting that its transition into an open, primed conformation promotes its lateral entrapment. Opening of SFKs triggers their catalytic activity because the SH2 and SH3 domains become accessible to interact with external ligands (Yadav & Miller, 2007), and the SH1 catalytic domain can be activated immediately after trans-autophosphorylation of Y420 (Young et al., 2001). In contrast, we found that the catalytic activity of the SH1 domain has little impact on the mobility of Fyn. This finding was demonstrated by the observations that (1) neither genetic or pharmacological suppression of Fyn activity had an impact on Fyn mobility, and (2) creation of an open-but-inactive Fyn through addition of the “kinase-dead” K299M mutant on top of the “open” Y531F mutant had little effect on Fyn immobilisation. Taken together, our findings indicate that the nanoscale organisation of Fyn is only affected by changes in its conformation, and that acquisition of an open, extended structure induces its lateral trapping in dendritic nanoclusters. In addition, as conformation and activity changes are interdependent in SFK enzymes, it is likely that the nanoclustering of Fyn facilitates its catalytic activity, being required to initiate its downstream signalling at the post-synapse.

The fact that SFKs are broadly expressed and share a common negative regulatory mechanism based on intramolecular interactions involving the SH2:tail and SH3:linker has motivated extensive research in exploiting these properties, to design strategies based on the creation of synthetic SH2/SH3-binding small molecules to finely control the activity of SFKs (Huang et al., 2016; Kukenshoner et al., 2017; Moroco et al., 2014; Yadav & Miller, 2007). Interestingly, monobodies (Huang et al., 2016; Huang et al., 2012) and peptoid-based ligands (Li & Lawrence, 2005) that are highly selective for the Fyn SH3 domain have already been generated to modulate the activity of Fyn kinase. We used a mEos2-tagged version of the G9 monobody created specifically to selectively bind Fyn SH3 domain (Huang et al., 2012). As Fyn interacts with tau through its SH2 and SH3 domains (Bhaskar et al., 2005; Lau et al., 2016; Lee et al., 1998), it is conceivable that binding of our G9-mEos2 monobody to the SH3 domain could induce a steric effect that interferes with the conformational mobility of Fyn in a similar way to tau. Our results indicate that binding of the G9-mEos2 monobody reduces the mobility of Fyn molecules, reinforcing the idea that alteration of their closed state by interfering with the intramolecular SH3:linker promotes an open-primed and immobilised Fyn conformation. The equilibrium dissociation constant (K_D_) for G9 monobodies bound to the Fyn SH3 domain calculated using isothermal titration calorimetry is 0.166 μM (Huang et al., 2012). In a different study using surface plasmon resonance, the K_D_ values calculated for WT tau and tau P301L bound to the SH3 domain of Fyn were 6.77 μM and 0.16 μM, respectively (Bhaskar et al., 2005). Although different techniques were used, these results suggest that G9 monobodies bind to the SH3 domain of Fyn with higher affinity than tau, and at comparable level as the FTLD mutant tau. These results are in line with our observations where overexpression of the mutant P301L tau decreased Fyn mobility to a similar extent as G9 monobodies do.

Interaction between tau and Fyn contribute to neurodegeneration associated with AD (Ittner et al., 2010) and FTLD-tau (Liu et al., 2020; Tang et al., 2020). Manipulating this interaction has been suggested as a potential therapeutical intervention. Decreasing either tau (Rapoport et al., 2002) or Fyn (Lambert et al., 1998; Um et al., 2012), for example, protects against Aβ toxicity in AD. However, undesired effects such as memory deficits associated with the absence of Fyn (Grant et al., 1992) suggest that further investigations are required to achieve more desirable therapeutic results. The recent development of a cell-permeable peptide inhibitor of the Fyn-tau interaction yielded promising results, reducing the endogenous Fyn-tau interaction and tau phosphorylation (Rush et al., 2020). G9 monobodies have also been used, demonstrating efficiency in inhibiting Fyn-tau binding (Cochran et al., 2014). Our findings on Fyn mobility using G9 monobodies against the SH3 domain are in line with these results and add the analysis of nanoscale mobility using super-resolution microscopy as a powerful new tool to evaluate potential therapeutical candidates.

Nanoclustering has been linked to the increased activity of multiple pre- and postsynaptic molecules, including AMPA receptors (Nair et al., 2013), syntaxin1A (Bademosi et al., 2017) and Munc18-1 (Chai et al., 2016). In particular, nanoclustering of syntaxin1A, which is associated with increased exocytosis, is driven by analogous mechanisms to those observed for Fyn. Syntaxin1A, a soluble N-ethylmaleimide-sensitive factor attachment protein receptor (SNARE) that drives vesicle fusion at the presynapse, switches from an inactive, closed conformation to an open conformation that can bind other SNARE proteins to initiate the fusion of synaptic vesicles with the plasma membrane. Interaction with the chaperone Munc18-1 promotes the folding of syntaxin1A into a closed and inactive state. Detachment of Munc18-1 facilitates the entry into an open conformation that allows syntaxin1A to interact with other SNARE proteins and form nanoclusters, thereby driving exocytosis at the presynapse (Kasula et al., 2016; Padmanabhan et al., 2020). Similarly, Fyn molecules fluctuate between a closed and an open conformation, the latter being enzymatically active and more immobile, and tau working as a molecular chaperone that modulates this transition.

We have previously reported that Fyn is organised in compacted nanodomains in dendrites (Padmanabhan et al., 2019), suggesting molecular crowding (Goose & Sansom, 2013; Li et al., 2016), spine geometry (Byrne et al., 2011) and interaction with neighbouring proteins as mechanisms that regulate the trapping and nanodomain organization of Fyn. Extension of Fyn molecules exposes their SH2 and SH3 motifs (Huculeci et al., 2016), and our results indicate that opening this conformation induces lateral trapping of Fyn into nanoclusters. However, it is unclear which interactions are responsible for this effect. Self-association has been described for other SFKs, with the formation of dimers enabling rapid potentiation of their activity when the enzymes adopt an open conformation (Irtegun et al., 2013). This suggests that multimerisation of opened, primed Fyn molecules could facilitate their clustering in dendrites. PSD95 is another possible candidate, as it stabilises molecules at the post-synapse and forms spine nanodomains of comparable size and frequency to those of Fyn (Hruska et al., 2018; Nair et al., 2013). Both PSD95 and Fyn associate with the plasma membrane following palmitoylation of specific residues (Sato et al., 2009; Tezuka et al., 1999; Topinka & Bredt, 1998) and PSD95 interacts with the SH2 domain of Fyn (Tezuka et al., 1999). This suggests that entry into an open conformation may induce Fyn to bind to PSD95, resulting in their postsynaptic nanoclustering. Tau also interacts with Fyn, and has previously been shown to control its localisation and nanoclustering in dendrites (Ittner et al., 2010; Padmanabhan et al., 2019). However, it is unclear whether this effect reflects a direct interaction with tau or is caused by tau’s regulation of the cytoskeleton in neurons. In this respect, tau interacts with actin (Cabrales Fontela et al., 2017; Elie et al., 2015) and the P301L mutant tau induces aberrant presynaptic actin polymerisation that is capable of crosslinking synaptic vesicles and restricting their mobilisation (Zhou et al., 2017). The actin cytoskeleton modulates nanoclustering of membrane proteins (Torreno-Pina et al., 2016), opening the possibility that it could also modulate the nanoscale organisation of Fyn. Further work is therefore needed to evaluate the potential involvement of the actin cytoskeleton in Fyn nanoclustering. Analysis of the kinetic parameters of the interaction between the SH3 domain of Fyn and pseudo-phosphorylated forms of tau and mutant forms associated with FTLD-tau, showed increased binding affinities when compared to wild-type tau, with the FTLD-tau mutant variant P301L having the highest SH3 affinity (Bhaskar et al., 2005). The interaction between Fyn and tau has been extensively characterised in terms of the residues responsible for this binding (Bhaskar et al., 2005; Cochran et al., 2014; Lau et al., 2016; Lee et al., 1998; Usardi et al., 2011; Wang et al., 2019) and the amino acids that are phosphorylated as a consequence (Lee et al., 2004). However, the crystal structure that would provide a detailed representation of how Fyn and tau interact is still missing. Although we lack precise structural information of Fyn-tau binding, it is therefore tempting to speculate that P301L tau may stabilize an open conformation, thereby promoting nanoclustering of Fyn at the post-synapse (Figure 8). Unfortunately, our temporal and spatial resolution limits do not allow us to determine whether the P301L mutant tau preferentially interacts with Fyn in its open state, or the binding of the mutant tau induces a conformational change that extends the structure of Fyn. Although it is possible that the nanoclustering of Fyn is facilitated by other interactions, our previous results suggest that P301L tau preferentially drives the cluster state of Fyn in hippocampal dendrites (Padmanabhan et al., 2019). Our new findings reveal the formation of a more immobile and active form of Fyn through interaction with P301L mutant tau, when compared to wild-type tau. Interestingly, in a FTLD-tau transgenic mice model, the loss of dendritic spines associated with impairment of spine plasticity occurs in the absence of detected hyperphosphorylated tau species or NFTs (Hoffmann et al., 2013). These findings are in line with observations from brain tissue of AD patients (Blazquez-Llorca et al., 2011; Merino-Serrais et al., 2013). Prefibrillar tau species such as tau monomers, dimers and higher order oligomeric aggregates are responsible for the neurotoxic effects in AD (Polanco et al., 2018), and have been suggested as a cause for the impairment in spine plasticity. However, our findings further suggest the neurotoxicity from P301L tau-mediated Fyn over-activity as a plausible complementary new mechanism responsible for the early loss in spine plasticity and morphology defects. Overall, the results presented in here support the idea that the ability of the tau to associate with Fyn is increased in FTLD-tau, altering the nanoscale organisation of Fyn and promoting an aberrant over-activity that potentiates neurodegeneration.

**Figure 8.**
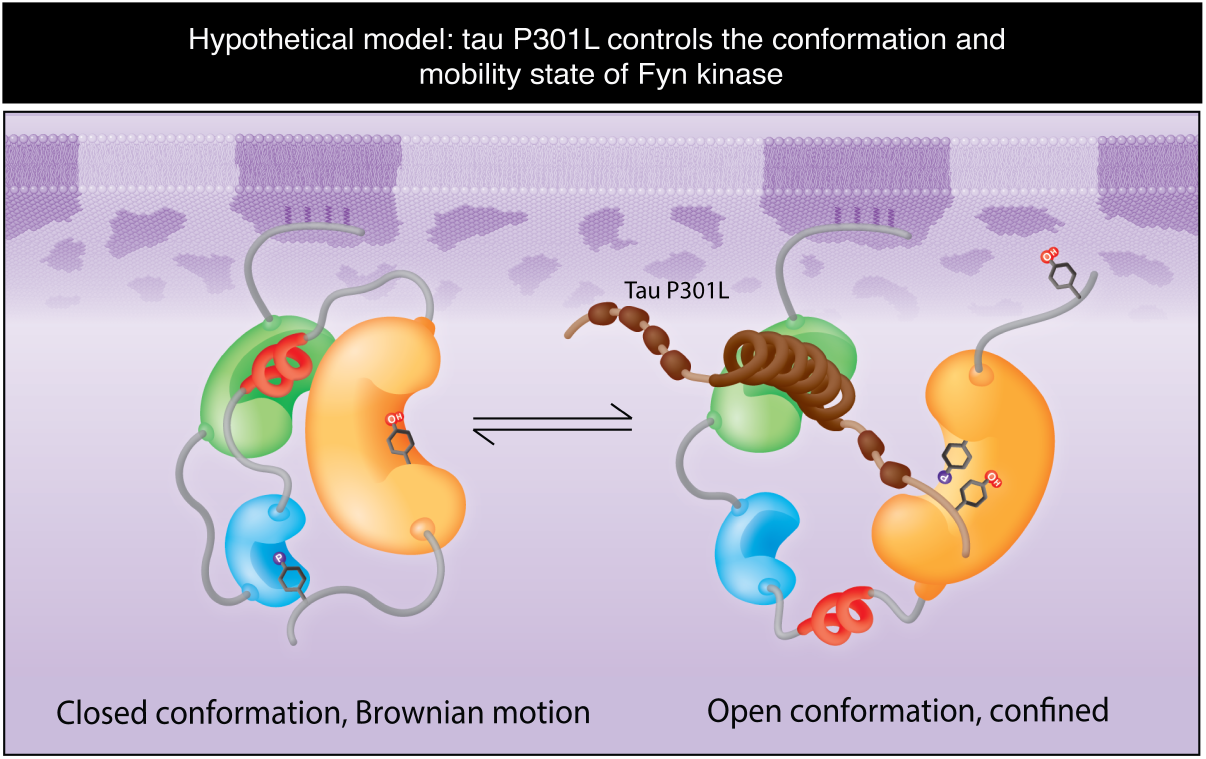
Model for the lateral entrapment of Fyn, mediated by entry into an open conformation. Tau P301L (brown) binds to the SH3 domain (green) of Fyn and displaces the interaction of SH3 with the linker region (red), which stabilises the open conformation of Fyn. The open conformation of Fyn bound to tau P301L exposes the SH2 domain (blue) of Fyn to interact with additional binding proteins, and positions tau P301L N-termini in close proximity with Fyn’s catalytic domain. This extended conformation also causes the lateral entrapment of Fyn.

## Experimental procedures

### Animal ethics and mouse strains

All experimental procedures were conducted under the guidelines of the Australian Code of Practice for the Care and Use of Animals for Scientific purposes and were approved by the University of Queensland Animal Ethics Committee (QBI/254/16/NHMRC). Wild-type mice (C57Bl/6 strain) used throughout the study were maintained on a 12-h light/dark cycle and housed in a PC2 facility with *ad libitum* access to food and water.

### Primary hippocampal cultures

Primary hippocampal neurons were prepared as described previously (Joensuu et al., 2017; Padmanabhan et al., 2019). Briefly, pregnant dams (C57Bl/6 mice) were euthanised using cervical dislocation, from which embryos (E16) were extracted and dissected in 1x Hank’s buffered salt solution (HBSS), 10 mM HEPES pH 7.3, 100 U/ml penicillin-100 μg/ml streptomycin (GIBCO-Thermo Fisher Scientific). Subsequently, the dissected hippocampal tissue was digested with trypsin (0.25% for 10 minutes). Digestion was halted by addition of Fetal Bovine Serum (FBS) (5%) (GIBCO-Thermo Fisher Scientific) with DNase I (Sigma) to prevent tissue clumping. The digested hippocampal tissue was left to incubate (37°C for 10 minutes). Following this, the hippocampal tissue was triturated and centrifuged (120 g, 7 min) and resuspended in neurobasal media, 100 U/ml penicillin-100 μg/ml streptomycin, 1x GlutaMAX supplement (GIBCO-Thermo Fisher Scientific), 1x B27 (GIBCO-Thermo Fisher Scientific) and Foetal Bovine Serum (FBS) (5%) (GIBCO-Thermo Fisher Scientific). Neurons were seeded in Poly-L-Lysine coated 29 mm glass-bottom dishes (Cellvis, CA, USA) at a density of 0.8-1 ×10^5^ neurons. A full media change was performed 2 hours after seeding using culturing media (Neurobasal Medium, 100 U/ml penicillin-100 μg/ml streptomycin, 1x GlutaMAX supplement, 1x B27).

### Heterologous cell cultures

HEK-293T cells were maintained in DMEM media (GIBCO-Thermo Fisher Scientific) supplemented with FBS (10%), 1x GlutaMAX and 100 U/ml penicillin-100 μg/ml streptomycin. Cells were transfected using the Lipofectamine™ LTX reagent according to the manufacturer’s instructions (Invitrogen-Thermo Fisher Scientific).

### Plasmids and reagents

The plasmid that codes for an FN3 monobody targeted against Fyn SH3 domain was a generous gift from Emeritus Professor Brian Kay (University of Illinois at Chicago, Illinois, US). The FN3 monobody was subcloned into the mEos2-N1 plasmid (Kasula et al., 2016) by inserting BamHI and EcoRI restriction sites at the N- and C-termini of the monobody, followed by BamHI/EcoRI digestion and ligation into mEos2 vector. Fyn mutant constructs were generated by site-directed mutagenesis, using the QuikChange II site-directed mutagenesis kit (Agilent), on the mEos2 donor vector containing full-length human Fyn isoform 1 (Padmanabhan et al., 2019) as a template. All the resulting plasmids were sequenced using the service of the AEGRC sequencing facility (The University of Queensland). PP2 and PP3 reagents were purchased from Calbiochem (Merck/Millipore).

### Western blotting

HEK293T cells were seeded in a 6 well plate and co-transfected with three combinations of plasmid (2 μg, 24 hrs with Opti-MEM and lipofectamine LTX with Plus Reagent): Fyn-mEos2 + GFP, Fyn-mEos2 + tau-GFP or Fyn-mEos2 + tau-P301L-GFP. HEK293T cells were isolated following trypsin digestion. Protein homogenates were boiled at 95°C in Laemelli buffer, run on a Tris-Glycine Precast Gel (4-15%, Bio-Rad) at 150V and subsequently transferred to a PVDF-FL membrane (Millipore), which was incubated in blocking buffer (Odyssey, TBS) for 1 hour at room temperature. Subsequently, membranes were incubated with primary antibody. Membranes were stained for total Fyn (1:1000 rabbit anti-Fyn, Cell Signalling Technology), active Src (1:1000 rabbit anti-phosphor-Y16, Santa Cruz), GAPDH (1:1000 mouse anti-GAPDH, abcam #189095) and GFP (1:1000 rabbit anti-GFP, Millipore #AB3080P); in blocking buffer overnight (4°C). Membranes went through TBS-T washes (5x) and incubated in secondary antibody (1:10000 donkey anti-mouse 680, Li-Cor Biosciences #32212 and donkey anti-rabbit 800, Li-Cor Biosciences #68023) with in Odyssey (TBS) blocking buffer (1 hour at room temperature). Fyn activity was measured as a ratio of phosphorylated Y416 intensity to total Fyn intensity, normalised to GAPDH.

### Single-particle tracking Photo-activated Localization Microscopy (sptPALM)

Neurons were transfected at DIV 14 using lipofectamine 2000 (ThermoFisher Scientific) (2 μg of GFP or mCardinal with 3 μg Fyn-mEos2 for 4 hrs) and were subsequently left to incubate for 24 hrs prior to imaging. Acquisitions (16 000 frames at 50Hz) were taken at 37°C on the Roper Scientific TIRF microscope fitted with an ILas^2^ double illuminator (Roper Scientific), a CFI Apo 100x/1.49N.A. oil-immersion objective (Nikon Instruments) and an evolve 512 Delta EMCCD cameras (Photometrics). A Perfect Focus System (Nikon) and an iLas2 double laser illuminator (Roper Scientific) was used for 360°C TIRF illumination. MetaMorph software (version 7.10.2, Molecular Devices) was used for image acquisition. A TIRF-quality ultra-flat quadruple beam splitter (ZT405/488/561/647rpc; Chroma Technology) for distortion-free reflection of lasers and QUAD emission filter (ZET405/488/561/640m; Chroma) were used. Transfected neurons were identified based on their GFP transfection using excitation with a 491-nm laser. For sptPALM, Fyn-mEos2 was photoactivated with low-levels 405-nm laser excitation (100 mW Vortran Laser Technology, 1-3% of initial laser power). Photoactivated Fyn-mEos2 molecules were subsequently photoconverted using 561-nm laser (150 mW Cobolt Jive, 70% of initial laser power).

### Data analysis

Tracking of single molecules of Fyn-mEos2 was performed in accordance with previous publications (Nair et al., 2013). Wavelet-based segmentation was used to detect single localizations of Fyn-mEos2. Following this, tracks of Fyn-mEos2 were computed using simulated annealing-based tracking algorithm implemented in PALM-Tracer, a software that operates in MetaMorph (Molecular Devices). Tracks lasting a minimum of eight frames were reconstructed and used to calculate the mean-square displacement (MSD) of Fyn-mEos2 using the equation MSD (t) = a + 4Dt, where D is the diffusion coefficient, a=y intercept and t=time. Trajectories with Log_10_[D] > -1.6 were considered to be mobile, allowing us to plot a frequency distribution histogram of Log_10_[D] and calculate the relative portion of mobile to immobile Fyn-mEos2 molecules (Bademosi et al., 2017; Constals et al., 2015). Fyn nanoclustering was quantified and visualised from sptPALM data using custom Python scripting based around the Density-Based Spatial Clustering of Applications with Noise (DBSCAN) functionality of the Python SciKit-Learn module (scikit-learn.org). Spatial centroids for each trajectory (with a minimum of eight steps) in a region of interest were clustered using DBSCAN with empirically determined “Goldilocks” values of ε = 0.1 µm (radius around each centroid to check for other centroids) and MinPts = 3 (minimum number of centroids within this radius to be considered a cluster). For each DBSCAN cluster, a convex hull of all the localizations comprising the clustered trajectories was used to determine the cluster area.

### Statistics

The D’Agostino and Pearson test was used to test for normality. For statistical analysis between two groups, a Student’s t-test was used. For multiple comparisons, a one-way ANOVA was used with the Tukey’s test for corrections for multiple comparisons. Statistical comparisons were performed on a per-cell basis (Fig 2-3, 5-7) and a per-cluster basis (Fig 4). The neurons analysed for each experiment were derived from a dissection of a minimum of five embryos. Unless otherwise stated, values are represented as the mean ± SEM. The tests used are indicated in the respective figure legends. For all statistical comparisons, p<0.05 was considered to be statistically significant. The tests used are indicated in the respective figure legends. Data were considered significant at p < 0.05. Statistical tests were performed, and figures were made using GraphPad Prism 7. A summary of statistical analyses is provided in Supplementary table 1.

## Figure legends

**Figure 2 – Figure supplement 1.**
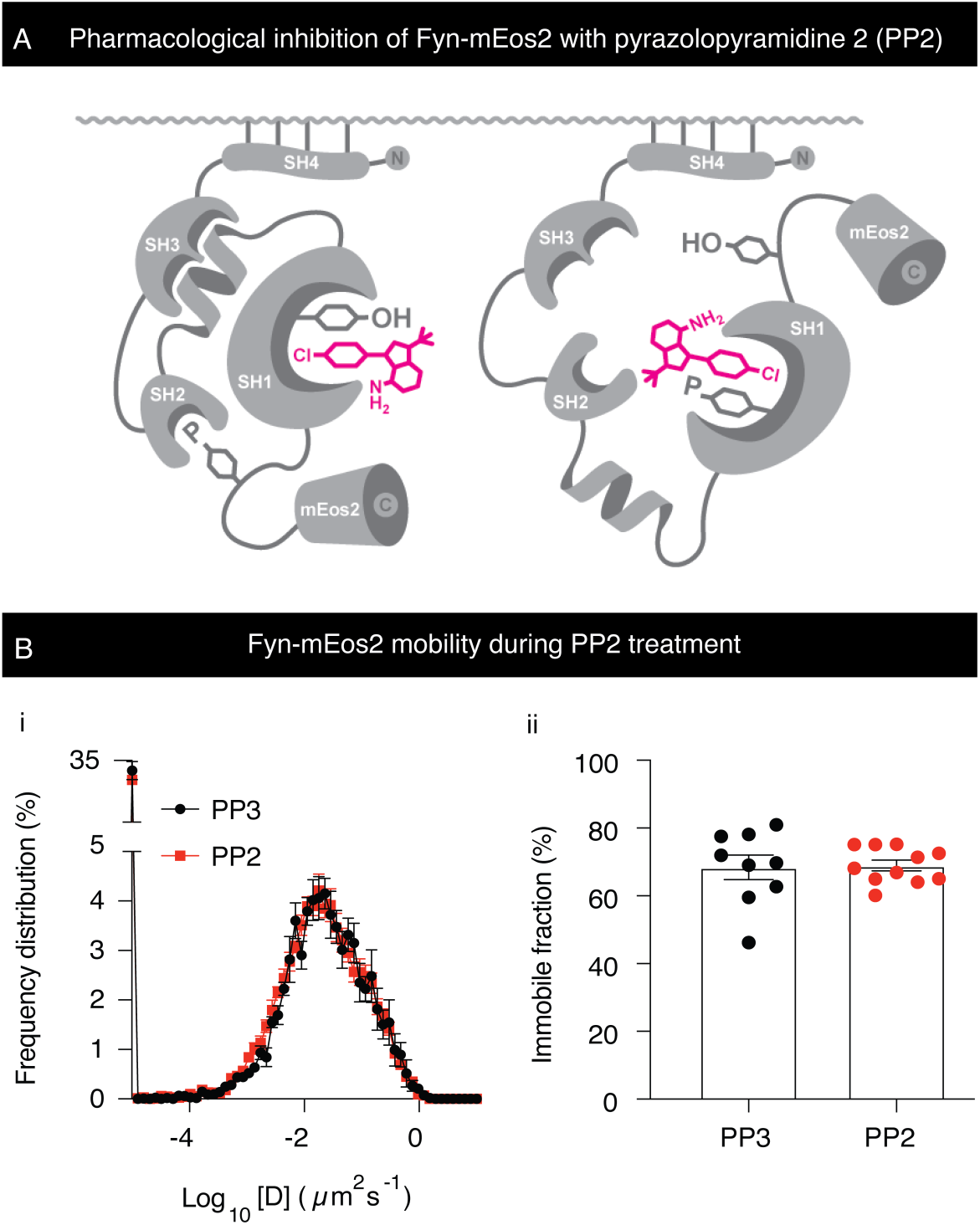
Pharmacological inhibition of the catalytic activity of Fyn with pyrazolopyrimidine 2 (PP2) does not impact its mobility. **A**. Illustration depicting the pharmacological inhibition of Fyn in closed and open conformation with PP2. **B**. Mobility of Fyn-mEos2. **(i)** Frequency distribution of Log_10_[D](µm^2^s^-1^) where [D] is the diffusion coefficient **(ii)** the immobile fraction (%). Error bars are standard errors of the mean (SEM). Mean ± SEM values were obtained for neurons transfected with Fyn-WT-mEos2 treated with PP3 (n=9) and PP2 (n=11). Statistical comparisons were performed using the Student’s t test.

## Article and author information

### Authors contribution

R.M.M. and F.A.M. were involved in conceptualisation of the project, design and supervision of the research. R.M.M. and C.S. performed super-resolution experiments. C.S. and R.M.M. analysed the data. T.W. contributed with analytic tools and analysis of the data. R.S.G. created specific plasmids. C.S., R.M.M., and F.A.M. wrote the paper. C.S., R.M.M, F.A.M and J.G. reviewed and edited the manuscript. F.A.M and J.G acquired required funding.

## Acknowledgments

We thank members of the Meunier laboratory for technical assistance, Nick Valmas for graphic design, Rowan Tweedale for the critical appraisal of the manuscript, Rumelo Amor and his team at the Queensland Brain Institute (QBI) Advanced Microscopy Facility for their excellent support with the microscopy, and Jean-Baptiste Sibarita (IINS, CNRS/University of Bordeaux) for his kind support and help with the single-molecule analysis software PALMtracer. We thank Brian Kay for generously gifting the FN3 monobody plasmids. R.M.M. was supported by The Clem Jones Foundation, The State Government of Queensland and the NHMRC Boosting Dementia Research Initiative. C.S was supported by a Research Training Program (RTP) scholarship. Single molecule imaging was performed at the Queensland Brain Institute’s Advanced Microscopy Facility, generously supported by the Australian Government through the ARC LIEF Grant (LE130100078 to FAM). This work was supported by the Federal Government of Australia (ACT900116) and the State Government of Queensland (DSITI, Department of Science, Information Technology and Innovation), and the National Health and Medical Research Council of Australia (GNT1145580 to J.G. and GNT1127999 to J.G. and F.A.M.) and a NHMRC Senior Research Fellowship (GNT1155794) to F.A.M.

